# Label-free amino acid identification for *de novo* protein sequencing via tRNA charging and current blockade in a nanopore

**DOI:** 10.1101/2020.06.25.170803

**Authors:** G. Sampath

## Abstract

A label-free procedure to identify single amino acids (AAs) for protein sequencing is developed in theory and simulated in part. A terminal AA cleaved from a protein/peptide, a tRNA, its cognate amino-acyl tRNA synthetase (AARS), and adenosine triphosphate (ATP) are brought together in a container where tRNA, if cognate, gets charged with AA and adenosine monophosphate (AMP) is released. The released AMP (and any free AA and ATP molecules) filters into the *cis* chamber of an adjoining electrolytic cell (e-cell) from where they pass through a nanopore into the *trans* chamber. Addition of NaOH to the container deacylates the tRNA if it is charged. The resulting free AA passes into the *cis* chamber of the e-cell, translocates into *trans*, and causes a current blockade; AA is immediately known from the identity of the tRNA (the two are cognate). If the tRNA is not charged there is no AA bound to it so AA remains unidentified. In this approach there is no need to distinguish among the 20 AAs by blockade size; it suffices to distinguish blockades from noise: thus a high-precision analog measurement has been transformed into a low-precision binary one. Identification is accurate because of tRNA superspecificity (the tRNA charging error rate is < 1/350); parallel execution with 20 different tRNAs can identify AA in one cycle. This is a *de novo* method in which no prior information about the protein is used or needed.

## 1. Introduction

Sequencing of a peptide or protein is commonly based on Edman degradation (ED) [1] followed by chromatography, on gel electrophoresis [2], or on mass spectrometry (MS) [3]; all three are bulk methods. The field is currently dominated by MS; for several decades more efficient methods have been sought, but success has been elusive. In contrast with DNA sequencing, which has progressed rapidly from Sanger sequencing to high throughput next generation methods that are highly automated, protein sequencing is yet to show similar progress. In large part this can be attributed to the fact that there are 20 amino acids (AAs) to identify, versus 4 bases in DNA. Many of them have similar physical and/or chemical properties, making it hard to tell them apart. Protein identification, in contrast, may be done without using any sequence knowledge or from a partial sequence. In this case affinity-based reagents may be used to identify a target protein [4]; this requires knowledge of the target. When identification is based on a partial sequence obtained with MS, the subsequence may be compared with a protein sequence database to determine if the subsequence occurs uniquely in some protein in the proteome [5].

The present work focuses on full *de novo* sequencing, no prior knowledge of the protein/peptide or information in a database is required.

Several new approaches to protein sequencing have emerged in recent years; for a recent review of protein sequencing research see [6]. In [7] a protein molecule is broken into short peptides and the carboxyl end of each peptide covalently attached to a glass slide. Fluorescent tags attached to selected residue types are used for detection at the N-end, following which N-end ED is used to cleave the terminal residue. Full *de novo* sequencing with this method is yet to be realized. A theoretical model is described in [8] in which peptides are assumed bound to a glass slide and the binding times of various optically tagged sensor molecules known as N-terminal AA binders (NAABs) [9] to the N-terminal residue of the peptide are calculated and the terminal residue identified. This method has the potential for full sequencing. Recently plasmonics, which is based on measuring changes in the refractive index of light due to an analyte, has been used to identify residues in a protein [10]. In [11] MD simulations are used to show that the protonated state of a terminal residue can be used to identify it.

Nanopores provide an electrically-based single molecule approach [12]. Unlike nanopore DNA sequencing, nanopore protein sequencing [13–15] is still in its infancy. A variety of methods are known, some in theory [16,17], others in practice [18,19] but yet to be developed fully. Recent related work has looked at recognizing single free AAs or residues within a peptide. In [20] 13 of the 20 standard AAs are identified from the volume they exclude as the AA translocates through a nanopore. In [21] naphthalene-based derivatives of 9 of the 20 AAs have been used as surrogates to increase their variability and allow easier identification with a nanopore. In [22] molecular dynamics (MD) simulations are used to show that blockades in a graphene nanopore can be combined with pulling forces in atomic force microscopy to identify AAs in a protein.

Most of the above methods are premised on precise measurement of analog quantities like blockade current, mass, electrical charge, fluorescence level, etc. The present work represents a departure from such a measurement-centered approach by using the ‘superspecificity’ property of tRNAs to correctly identify all 20 AAs. The measurement component is limited to determining whether a current blockade occurs in a nanopore or not (the exact size of the blockade is not relevant). This converts a high-precision analog measurement problem to a minimalist low-precision binary measurement problem.

### 1.1 The present work

This communication describes in theory, and simulates in part, a method for AA identification that is aimed at *de novo* protein sequencing for use in the bulk or with a few molecules. It combines chemical-based methods with nanopore-based sensing. At its center is the ‘superspecificity’ [23] property of transfer RNAs (tRNAs), which is used by living cells to translate mRNA to protein [24]. Thus for every AA there is a tRNA that binds to that AA and to no other; the in vitro error rate is about 1 in 350 [25]. Binding is accompanied by the release of adenosine monophosphate (AMP).

There are four steps: 1) confinement of reactants (AA, tRNA and its cognate amino-acyl tRNA synthetase (AARS), and adenosine triphosphate (ATP)) in a cavity, which results in charging of the tRNA with AA if cognate and release of adenosine monophosphate (AMP); 2) filtering or separation of tRNA and AARS from ATP, AMP, and AA; 3) deacylation of the tRNA, resulting in the release of AA if tRNA is charged; and 4) translocating the released AA through a nanopore in an e-cell. This unambiguously identifies AA from the identity of the tRNA. When executed in parallel on 20 or more copies of AA each with a different tRNA, the matching AARS, and ATP, AA is identified in a single cycle. By feeding terminal AAs cleaved from a peptide/protein to this parallel processor, *de novo* sequencing can be done.

## 2. A four-step procedure to identify an amino acid

This method is centered on the charging of a tRNA molecule with a cognate AA. It may be done in the bulk or with a few molecules. The following equations summarize the charging process [24]:

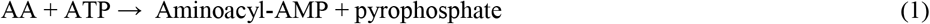

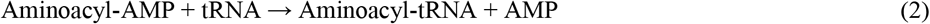

The procedure consists of four steps, corresponding to (a,b), (c), (d/f) or (g/h) in Figure 1, where / distinguishes between two cases: charged and uncharged tRNA.

**Fig. 1.**
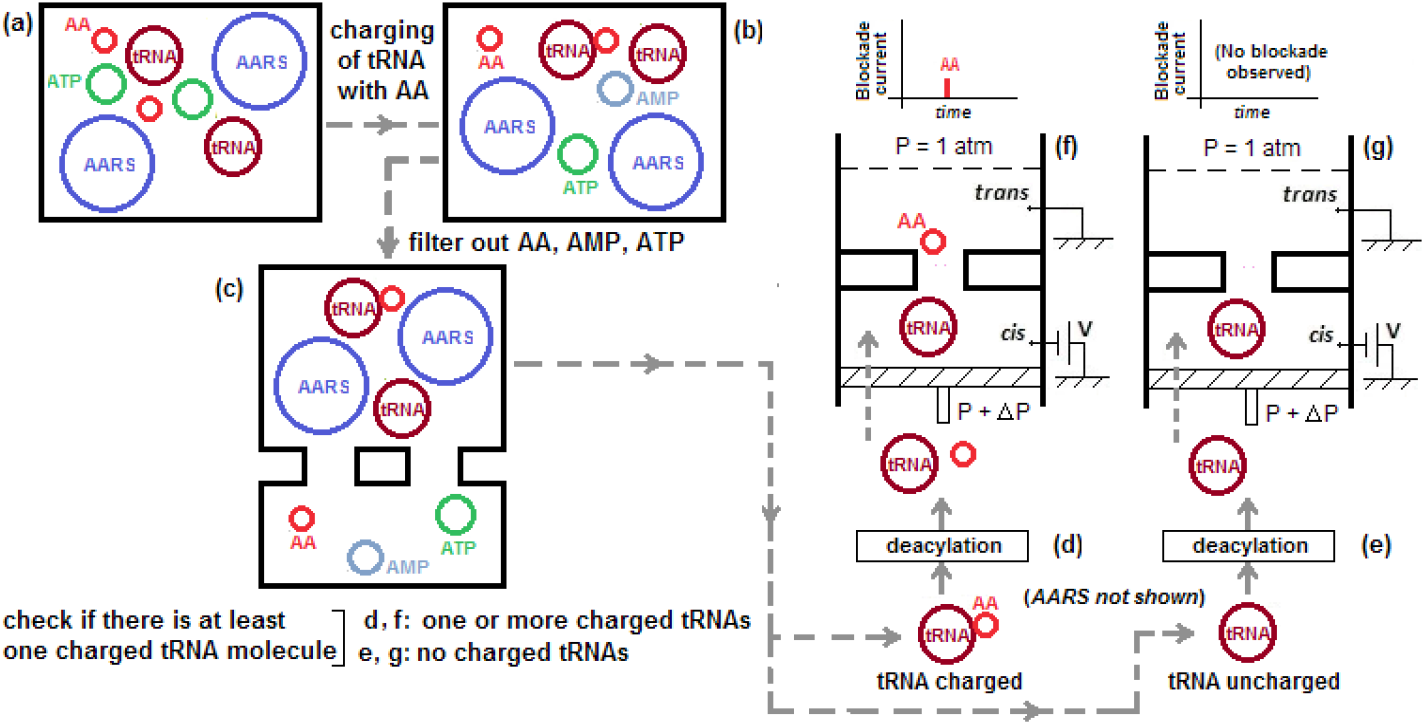
Amino acid identification process. (a,b) Charging of tRNAi (0 ≤ i ≤ N), one of N (20 ≤ N ≤ 61) different tRNAs; (c) separation of tRNAi, charged or not; (d) deacylation of charged tRNAi, (e) deacylation of uncharged tRNAi has no effect, products transferred to *cis* chamber of e-cell; (f) If tRNAi is charged free AA causes blockade → AA identified from tRNAi; (g) if tRNAi not charged, no blockade → AA not identified. Schematic, not to scale.

### Step 1: Charging of tRNA

The reactants need to be restricted to a small space because otherwise diffusion takes over. Such restricted spaces may take the form of sub-micro- to nano-volume containers or cavities. With a reservoir connected to a cavity, polymer molecules from the former can be trapped inside the latter. In [26] a pore-cavity-pore structure was introduced for the study of single molecules and time-of-flight measurements. This was modified in [27] for the analysis of long DNA strands. A femtoliter-sized ‘cage’ is used to trap a DNA strand that has passed through a nanopore into the cage from a source chamber on the other side of the pore. The trapped DNA is fragmented in the cage by a restriction endonuclease that enters the cage from the reservoir. The fragments are allowed to translocate through the pore back into the source chamber and the resulting current blockades used to study the fragments.

Figure 2 shows how molecules of AA, tRNA, AARS, and ATP are brought together in solution and kept in close proximity without diffusing away by hydraulic pressure [28], which works with all the analyte types involved, electrically charged or not. Simulations of a simple hydrodynamic model based on Poiseuille flow show that the reactants are confined to the ‘bottom’ of the container; see below. Note that no measurements are made on the reservoir-cavity structure so the dimensions do not have to be exact. The dimensions of the reservoir, passage, and cavity are about the same order as those in [26,27]. The simulation details are given in a Supplement.

**Fig. 2.**
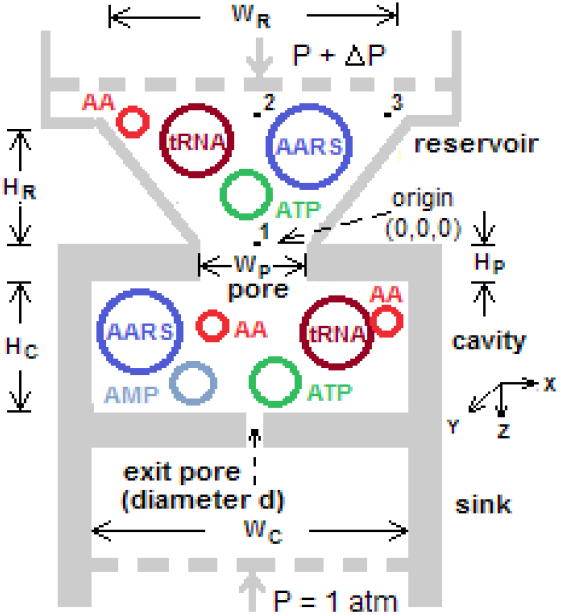
Schematic (not to scale) of reservoir-cavity structure showing confinement of reactants in cavity under hydraulic pressure and charging of tRNA with cognate AA. tRNA. AARS, and ATP molecules (Equations 1 and 2) are added to buffer solution containing AA molecules in reservoir. Taper in reservoir ensures that particle enters connecting pore and translocates into cavity. Hydraulic pressure P_atm_ + ΔP (where P_atm_ = atmospheric pressure and ΔP is the applied pressure) provides bias in z direction toward cavity bottom. H_R_ = height of reservoir; W_R_ = width of reservoir; W_P_ = width of pore connecting reservoir and cavity; H_C_ = height of cavity; W_C_ = width of cavity (see Supplement for values used in simulation). 1, 2, 3: Different origins for particle. Exit pore diameter d = 3-6 nm allows separation of smaller molecules from tRNA and AARS.

### Step 2: Separation of tRNA (charged or uncharged)

After a suitable incubation time in the container (to be determined experimentally) the products may include charged cognate tRNA, uncharged tRNA, free AA, AARS, ATP, and AMP. The last will be present only if tRNA is cognate and at least one tRNA molecule has been charged. tRNA, charged or not, and AARS are separated from AA, ATP, and AMP (if any); next the tRNA is separated from AARS (Figure 1d). The filter sizes required are calculated from the axes of enclosing ellipsoids for each of the analytes involved. Table 1 is a subset of Table S-1 in the Supplement; it shows the axis sizes of the covering ellipsoids for each type of reactant. Referring to Table 1, a 3-6 nm diameter pore can separate tRNA and AARS from AA, AMP, and ATP; and a 9-10 nm diameter pore can separate tRNA from AARS.

**Table 1.** Sizes of reactants involved in tRNA charging. Amino acids represented by smallest (Glycine) and largest (Tryptophan).

| Reactant | Axes of ellipsoid enclosing reactant |  |  |
| --- | --- | --- | --- |
|  | Major (nm) | Minor (nm) | Minor (nm) |
| Glycine (AA) <sup>1</sup> | 0.56 | 0.35 | 0.21 |
| Tryptophan (AA) <sup>2</sup> | 1.10 | 0.7 | 0.34 |
| AMP <sup>3</sup> | 1.67 | 0.95 | 0.4 |
| ATP <sup>4</sup> | 2.34 | 1.09 | 0.44 |
| Phe-tRNA <sup>5</sup> | 10.56 | 6.54 | 3.64 |
| Ala-AARS <sup>6</sup> | 13.94 | 9.41 | 8.46 |

<sup>1,2,3,4,6</sup> Atomic coordinate data from files.rcsb.org/ligands/view/AMP\_ideal.sdf, files.rcsb.org/ligands/view/ATP\_ideal.sdf, www.rcsb.org/ligand/GLY, files.rcsb.org/ligands/view/TRP\_ideal.sdf, www.rcsb.org/structure/1YFS. (The last is for Ala-AARS.)
<sup>5</sup> Atomic coordinate data from [29].

However it may not be necessary to separate tRNA from AARS (see note at the end of Step 3). In this case the exit pore at the bottom of the cavity can act as a filter to separate the the smaller molecules (AA, AMP, ATP) from tRNA and AARS, which remain in the cavity; see Figure 3 below, which is an extension of Figure 2 to which an e-cell has been adjoined, with the sink being replaced by the *cis* chamber of the e-cell. Figure 3a shows the filtration effect when there is a charged tRNA in the cavity. In this case, charging would have caused an AMP to be released (see Equation 2 above), the AMP exits the cavity and translocates through the nanopore in the e-cell into the trans chamber of the e-cell. Figure 3b shows the effect when there is only an uncharged tRNA in the cavity, notice the absence of an AMP molecule in the filtered output from the exit pore. (The bottom part of the figure shows blockades caused by the filtered-out ATP, AMP and AA as they translocate through the the nanopore of the e-cell. The AMP blockade in Figure 3a can be used to identify AA, see Section 3.)

**Fig. 3.**
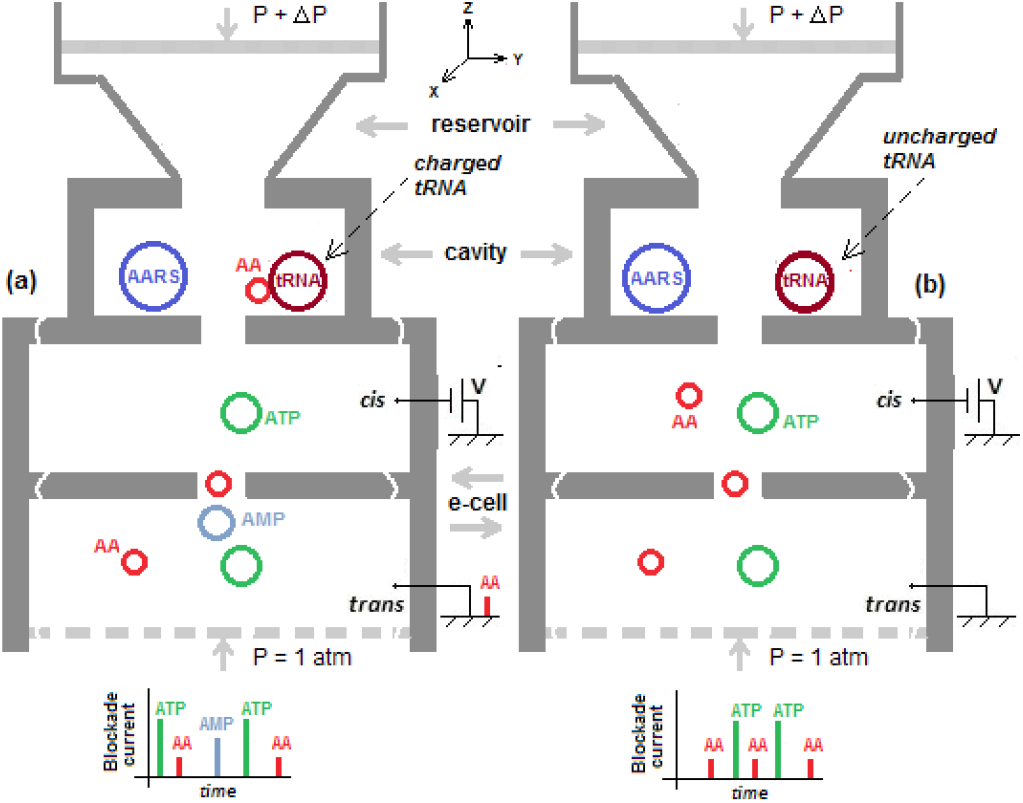
Integrated device for amino acid identification extends reservoir-cavity structure with electrolytic cell (e-cell) replacing sink. AA, ATP, and AMP in cavity have filtered through connecting pore at bottom of cavity into cis chamber of e-cell, then translocate through nanopore of e-cell into trans chamber. (a) Cavity has charged tRNA (brown) with AA (red) bound to it. Free AA, AMP (blue) released by charging of tRNA, and free ATP shown translocating into trans. (b) Cavity has uncharged tRNA (brown, no AA attached). Free AA and free ATP (green) shown translocating into trans. (Note absence of AMP molecule.) Hydraulic pressure applied to reservoir at top. (Bottom of figure shows blockades due to translocation of AA, ATP, and AMP through e-cell’s nanopore. AMP blockades can be used for AA identification, see Section 3.)

### Step 3: Deacylation of charged tRNA

The tRNA, if charged, can be deacylated to release the bound AA, which can then be detected in the nanopore of an e-cell. Deacylation can be done enzymatically or without an enzyme. Non-enzymatic deacylation is much simpler. It may be done with NaOH or by controlling the solvent’s pH level [30]. The products of this step are either tRNA and the dissociated AA (if tRNA was charged) or only tRNA. If it is assumed that deacylation has no effect on AARS then there is no need to separate tRNA from AARS and only one filtration step is required with a 3-6 nm pore. Deacylation can be achieved by adding NaOH to the reservoir in Figure 3 after an empirically determined time for confinement of reactants (Step 1) and filtration (Step 2) to complete. In the latter case all the free reactants have been translocated fully into the trans chamber of the e-cell, and the cavity contains only AARS and charged or uncharged tRNA.

### Step 4: Identification of AA from current blockade

Deacylation releases a bound AA from a charged tRNA, the free AA then filters through the exit pore at the bottom of the cavity into the cis chamber of the e-cell and translocates through the e-cell’s nanopore to cause a blockade. If there is no charged tRNA in the cavity, no such blockade would be observed. Figure 4a shows the former, Figure 4b shows the latter.

**Fig. 4.**
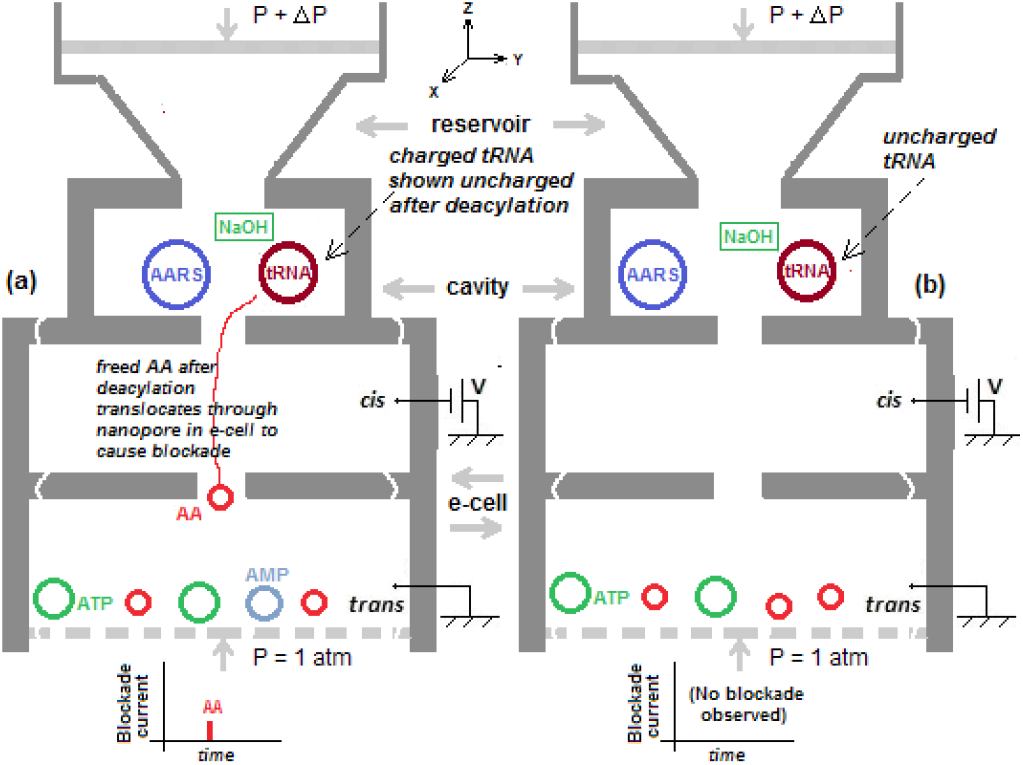
Result of introducing NaOH into cavity from reservoir (Step 3). (a) NaOH deacylates charged tRNA; freed AA translocates through nanopore of e-cell to cause blockade; (b) tRNA is uncharged, NaOH has no effect; no free AA, no blockade. (Molecules at bottom of trans chamber are remnants from preceding filtration step.)

In the first case AA is immediately known from the identity of the tRNA. The only requirement is that the blockade level due to the smallest volume AA, Glycine (G), be distinguishable from baseline noise. This is discussed later in the context of nanopore noise.

The above leads to the following simple determination:

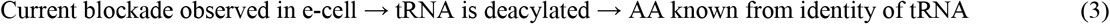

Conversely,

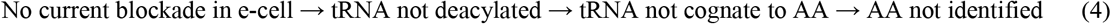

Note that the tRNA that enters the e-cell at the beginning of Step 4 does not contribute anything to the blockade measurement (it is too big to translocate through the pore). Only the dissociated AA (Ser in the example) is involved.

Assuming that the size of the current blockade I_B_ in the e-cell is roughly proportional to V_AN_, the volume excluded when an analyte AN passes through the pore [19,20] can be written as

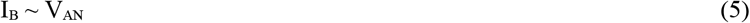

The observed exclusion volume due to AA satisfies

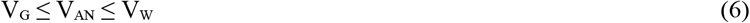

where V_G_ is the volume of the smallest AA, Glycine (G), and V_W_ that of the largest AA, Tryptophan (W). From Table S-4 in the Supplement, V_G_ = 59.9 × 10^−3^ nm^3^ and V_W_ = 227.9 × 10^−3^ nm^3^. (Equation (6) is an approximation, see Item 3 in Section 4 for a related brief discussion.)

#### Feasibility and accuracy: conditions for correct identification of an amino acid

The feasibility of the method described above can be judged from the feasibility of the parts: tRNA charging; AMP release; nanopore sensing of AA blockades; hydraulic pressure in a nanopore; non-enzymatic deacylation; and cleaving of a terminal residue from a protein. All of these are based on known and well-established laboratory techniques.

The accuracy of the method can be judged based on a set of necessary conditions for correct identification of an AA. The following assumptions are made:

1. Superspecificity holds in vitro. The considerable literature on in vitro experiments involving AARSs supports this assumption; for a recent brief review see [31].
2. With enough reactant molecules (AARS, ATP, AA) and favorable environmental conditions (temperature, buffer, etc.) at least one tRNA molecule gets charged. The results of in vitro experiments on lysyl-tRNA described in [32], with product lifetimes in the tens of minutes, make this a reasonable assumption.
3. Buffers and other reagents (such as NaOH for deacylation) do not confound observation of blockades due to any of the 20 AAs. This is supported by the extensive literature on blockade measurements in nanopores.

Four conditions have to be satisfied in order for the above procedure to correctly identify an AA (and do so in a reasonable amount of time):

1. The reactants necessary for charging of a tRNA must be in close proximity until successful charging occurs.
2. tRNA molecules are perfectly separated from the other reactants; filtration times must not be excessive.
3. Deacylation of a charged tRNA always occurs.
4. Current blockades due to any AA are distinguishable from baseline noise.

##### Condition 1

With bulk samples, this condition is easily achieved. With trace amounts of AA, the available AA molecules must be confined in a micron- or sub-micron-sized cavity with ATP and a tRNA and its related AARS to counter the dispersive effects of diffusion. AMP, ATP, and tRNAs carry a negative charge so an electrophoretic field can be used, but this is not always the case with AAs and AARS. Thus 7 of the 20 standard AAs carry a positive or negative electrical charge whose value depends on the pH of the solution, the other 13 are neutral [33]. An AARS is a protein with a net positive, negative, or zero charge whose value is determined by the primary sequence. The solution is to use a charge-independent force field. An example is hydraulic pressure, which has been used in the study of polymer translocation through nanopores [28]. Thus while diffusion is isotropic the directed hydraulic force biases the movement of particles toward the bottom of the cavity thus ensuring that they accumulate there. This process was simulated with Monte-Carlo techniques by assuming an analyte molecule to be a dimensionless particle. Figure 2 shows a schematic of the reservoir-cavity structure simulated. The simulation procedure is similar to the procedures used in [34], which is partly based on [35]. In the latter the terminal residue of a DNA strand passing through the first pore of a tandem electrolytic cell is cleaved by an exonuclease on the exit side of the pore, enters and translocates through a second nanopore, and is identified by the level of the blockade caused in the second pore. See the Supplement for a description of the geometry, protocol, and computations involved.

To assess the efficacy of confinement the reservoir-cavity structure is simulated with a range of parameters (diffusion coefficient, hydraulic pressure level, origin of particle, and number of diffusion steps). The following measures are used: 1) Time of entry of particle into cavity; 2) Time to reach bottom of cavity; and 3) Dwell time at bottom of cavity. The simulation is stopped when the dwell time exceeds 106 continuous steps. In all runs the particle was found to always settle at the bottom of the cavity. Table 2 shows the results. The results suggest that if the simulation were to continue the particle would remain indefinitely near the bottom of the cavity. This of course is the desired outcome.(This may be compared with the results obtained in [26,27], where similar structures are used; confinement times range from a few milliseconds [26] to tens of minutes [27].) The lower part of Table 2 shows the results for when the motive force is exclusively diffusive (no hydraulic pressure). The results show that the particle is still diffusing in the reservoir at the end of the run. Hydraulic pressure is key to confinement of the reactants to the cavity.

**Table 2.** Simulation of dimensionless particle in reservoir-cavity structure under hydraulic pressure (= 1 atm) and diffusion

| D | Origin | $T_1 (\times 10^{-6} \text{ s})$ | $T_2 (\times 10^{-6} \text{ s})$ | $T_3 (\times 10^{-6} \text{ s})$ |
| --- | --- | --- | --- | --- |
| 0.3 | (0,0,0) | 1.21 | 1.38 | > 1.0 |
|  | (0,0,-H<sub>R</sub>) | 1.53 | 1.70 | > 1.0 |
|  | (W<sub>R</sub>/2, W<sub>R</sub>/2, -H<sub>R</sub>) | 1.62 | 1.79 | > 1.0 |
| 3.0 | (0,0,0) | 1.23 | 1.40 | > 1.0 |
|  | (0,0,-H<sub>R</sub>) | 1.47 | 1.64 | > 1.0 |
|  | (W<sub>R</sub>/2, W<sub>R</sub>/2, -H<sub>R</sub>) | 1.62 | 1.79 | > 1.0 |
| 30.0 | (0,0,0) | 1.18 | 1.34 | > 1.0 |
|  | (0,0,-H<sub>R</sub>) | 1.25 | 1.41 | > 1.0 |
|  | (W<sub>R</sub>/2, W<sub>R</sub>/2, -H<sub>R</sub>) | 1.63 | 1.80 | > 1.0 |
| Pure diffusion |  |  |  |  |

| Origin | $N_4$<br>($\times 10^8$) | $z$<br>($\times 10^{-6}$ m) | $z_{\max}$<br>($\times 10^{-6}$ m) | Entered<br>cavity? | Reached cavity<br>bottom? |
| --- | --- | --- | --- | --- | --- |
| (0,0,0) | 4 | -8.82 | 0.0059 | Yes | No |
| (0,0,-$H_R$) | 4 | -9.75 | -9.6616 | No | No |
| ($W_R/2, W_R/2, -W_R$) | 4 | -9.90 | -9.6722 | No | No |

##### Condition 2

Referring to Table 1 tRNA is clearly separated in size from AARS on one side, and from AMP, ATP, and any of the AAs on the other. Thus filtration can be done with nanopores of appropriate size. Assuming that the addition of NaOH in Step 3 does not adversely affect AARS, it is not necessary to separate tRNA from AARS, so a single filtration step to separate tRNA and AARS from AMP, ATP, and AA is sufficient.

Filtration times must be reasonable. The following calculations show that this condition can be met if the exit pore in Figure 2 has a diameter in the range 3-6 nm.

Consider for simplicity a cubical cavity of side 5 μm (volume = 125 × 10^−18^ m^3^ = 125 femtoliters). Consider a nanopore with a diameter of 5 nm (which will allow AMP, ATP, and any of the 20 AAs to filter through but not Phe-tRNA or phenylalanyl-tRNA synthetase) and a thickness of 5 nm. A solid-state pore with these dimensions is possible. With Poiseuille flow the flow rate is Q = πR^4^ΔP/8ηL, where R = radius of pore = 2.5 × 10^−9^ m, L = thickness of pore = 5 × 10^−9^ m, ΔP = pressure difference = 2 atm = 2 × 10^5^ Pa, η = solvent (water) viscosity = 0.001 Pa.sec. This leads to

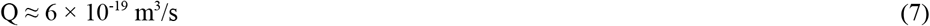

The time required to evacuate the cavity is

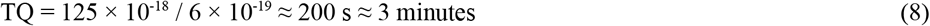

Compare this with a biological nanopore, such as α-Hemolysin (AHL). The barrel in AHL has a diameter of 2.6 nm and a length of 5 nm. In this case the filtration time for the same cavity is about 45.6 mins, and about 27 s with a 10 × 10 array.

##### Condition 3

In the living cell deacylation is done by an enzyme to correct charging errors [25]. Thus if a tRNA is misacylated an editing AARS deacylates the tRNA and attaches the correct AA to it. A table of Class I and Class II AARSs and their AA editing targets is given in [31]). Here non-enzymatic deacylation with NaOH is assumed; see [30] for a description of its use in the deacylation of cysteine-tRNA.

##### Condition 4

When AA is cognate to tRNA it is deacylated in Step 3, enters the e-cell with tRNA as a free AA, and causes a blockade in the e-cell. Let the quiescent or base pore current, that is, the current through the pore in the absence of an analyte, be I_base_. Considering Equation (6), I_base_ ~ V_pore_ = volume of pore = πd^2^h/4, where d is the diameter of the pore and h its thickness. Let the blockade size or drop in the base current due to AA be IB_AA_. The smallest current drop IB_min_ is due to the AA with the smallest volume, namely Glycine, G. IB_min_ = I_B-G_ ~ V_AA_=G = 0.059 nm^3^ (from Table S-4 in the Supplement).

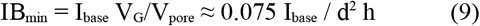

with d and h in nm. With a base current of 100 pA, d = 1 nm, and h = 2 nm, IB_min_ = 2.5 pA.

IB_min_ must be distinguishable from the baseline noise current I_RMS_, which is the rms (root-mean-square) value of the noise around Ibase. Define the signal-to-noise ratio (SNR) for analyte AN as

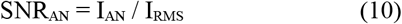

where I_AN_ is the size of the current blockade due to AN. A sufficiently high SNR can ensure that the blockade current due to Glycine can always be distinguished from I_RMS_. Figure 5 shows the probability that a measured blockade for Glycine is < Th_noise_ = 2 I_RMS_ for different values of σ_AA=G_. Here Th_noise_ is the volume threshold for the current noise I_RMS_ amd σ_AA_ is the standard deviation of the volume of AA. (Volume information for the 20 AAs is given in Table S-4 in the Supplement.)

**Fig. 5.**
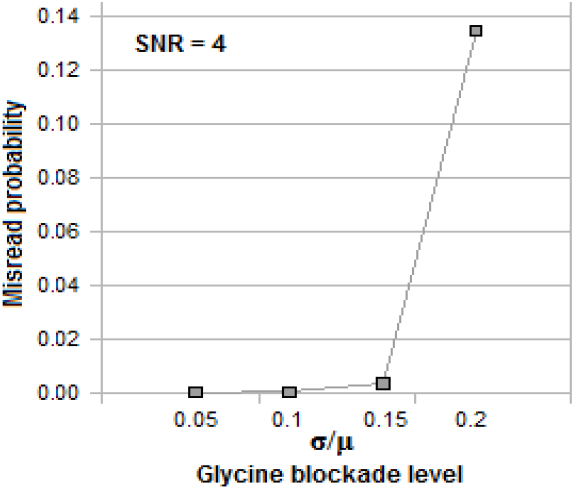
Probability of 2*baseline RMS noise (I_RMS_) in nanopore current being mistaken for measured blockade due to smallest volume AA glycine (G) with a signal-to-noise ratio (SNR) of 4 for different values of standard deviation σ relative to mean volume μ of G.

Various methods of increasing the SNR for both kinds of pores are discussed in [36]. In general biological pores have higher SNRs than solid state pores. The latter have SNRs in the range 3-40, which is adequate for the present purpose. Additionally slowing down the analyte as it translocates through the pore can lead to a significant increase in SNR by reducing the bandwidth required for detection and at the same time decreases high frequency noise; see Section 4.

#### Parallel processing of N copies of AA with N different tRNAs

The sequence of steps given above can be done in parallel with N different tRNAs and N copies of AA, filters, and e-cells, as shown in Figure 4. With N = 20 superspecificity of tRNA guarantees that AA is identified by one of the 20 units.

Figure S-4 in the Supplement shows a trace of the identification procedure through the 20 units with AA = Ser. With this parallel processor the identity of AA is immediately known from the number of the unit in which a blockade is observed in the e-cell. No post-processing or calculations of any kind are required. Importantly, in the downstream e-cell that contains the free AA there is no need to know the exact size of the blockade. Unlike in other nanopore sequencing methods there is no need to discriminate among the 20 AAs when measuring blockades.

Notice that the only measurement performed is for determining whether there is a blockade or not. This is a simple binary measurement: In 19 e-cells the answer is No, in exactly one e-cell the answer is Yes.

In fact each of the 20 e-cells is uniquely tied to one of the 20 AAs, no analyte other than that AA can enter the pore in the e-cell in Step 4. This makes possible design of each of the 20 units for optimal performance that is tailored to the AA associated with that unit. This is discussed in Section 4 under ‘Silo effect’.

#### Processing in series if only a few molecules are available

If only a few molecules of AA are available (as happens if the sample comes from single cells) the processing can be done in series, as shown in Figure 7. Pipelining speeds are determined by the slowest stage in the pipeline. An in-depth study of serial processing of AAs for identification based on the procedures given here remains to be done.

**Fig. 6.**
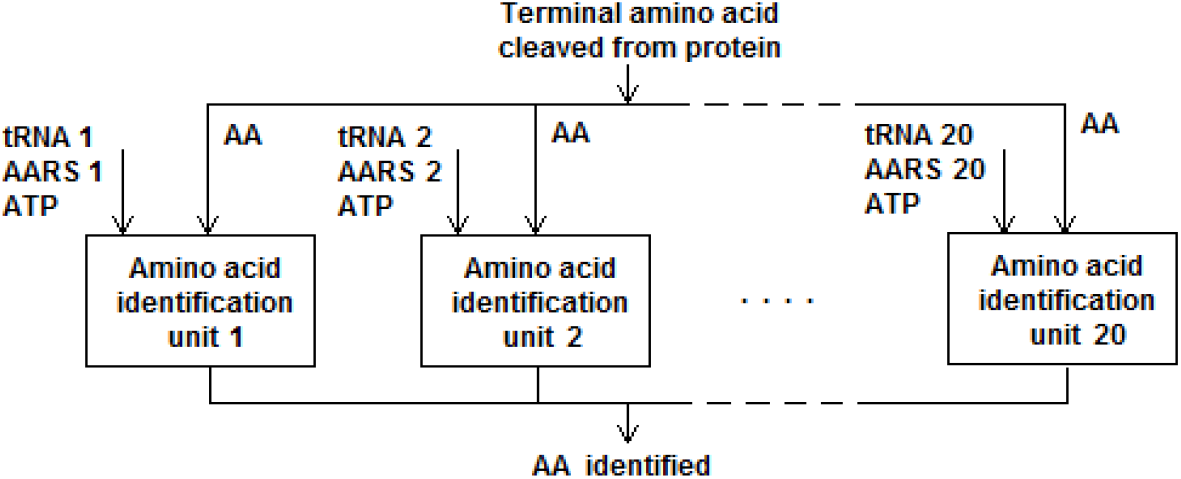
Parallel architecture for single AA identification

**Fig. 7.**
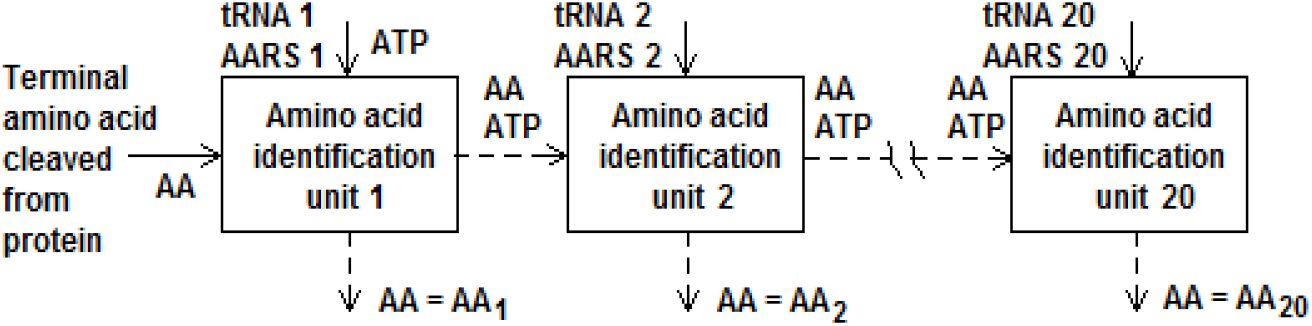
Series architecture for single AA identification

#### Probability of correct identification

The probability that AA is correctly identified can be estimated by considering the number of molecules of each reactant in the cavity in Step 1 and the number of charged tRNAs that enter Step 3 for deacylation. Assuming no charging errors, let 1 of M_1_ tRNA molecules get charged in the cavity and 1 of M_2_ charged tRNA molecules be deacylated. In a parallel implementation with 20 units, a total of 20 M_1_M_2_ AA molecules in the sample and 20 M_1_M_2_ tRNA molecules can identify AA with probability 1. With M_1_ = M_2_ = 10 (corresponding to charging and deacylation success rates of 10%), this is about 3 zeptomoles, which is comparable to the molar volumes of samples used in a recent label-based protein sequencing method [7].

#### Protein sequencing

The full peptide or protein sequence can be obtained by applying the above procedure to terminal residues cleaved in succession from a peptide/protein. Cleaving may be done with ED [1], which is known to be a harsh process and is not suited to the present approach because ED produces derivatized AAs. Less harsh alternatives that produce free AAs, such as carboxyl peptidases [37] and aminopeptidases [38] are more appropriate; see [16] for a brief discussion. In [39] an enzyme designed on a computer and named Edmanase that cleaves residues at the N-end of a protein is described.

## 3. AA identification from AMP blockades

AA can also be identified from the AMP molecule that is released during the charging process. However the blockade caused by AMP as it translocates through the nanopore of the e-cell must be distinguished from blockades due to some of the other reactants (ATP, free AAs) translocating through the pore. While this introduces some error, the error rate is still < 10%; this is discussed below.

### Error analysis

The above identification process based on AMP blockades can be analyzed for potential false negatives and false positives. A false negative occurs if the tRNA is charged and ATP were to cause an AA-sized blockade, or the ATP looks like an AMP and the AMP looks like an ATP. It is sufficient to consider the former possibility as it has the higher probability. A false positive occurs if the tRNA is not charged and AA were to cause an AMP-sized blockade. Figures 8a and 8b show the variation of the corresponding two probabilities PFN and PFP for each of the 20 AAs for four values of σ (0.05μ, 0.1μ, 0.15μ, 0.2μ), where μ and σ are the mean and standard deviation of a normal distribution of analyte volume (see Table S-4 in the Supplement). The worst case occurs with W (Tryptophan), the largest volume AA, with a 17% false negative probability when σ = 0.2μ. If the two types of errors are averaged over all 20 AAs weighted by their frequency of occurrence in the human proteome (Uniprot Id UP000005640; 11326153 residues) the weighted error probability is 9.5% for σ = 0.2μ; see Figure 8c. The details can be found in the Supplement.

**Fig. 8.**
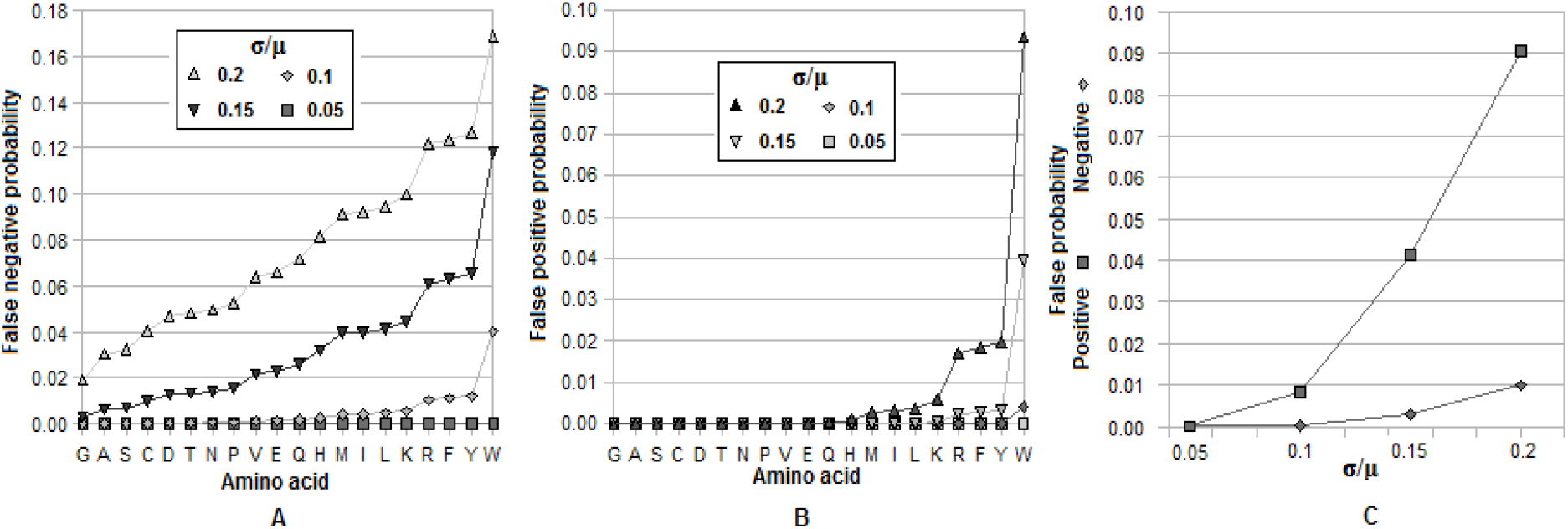
Error probabilities for the standard 20 amino acids. (a) False negatives; (b) False positives; (c) Both over the full set of proteins in the human proteome weighted by frequency of occurrence of each amino acid.

## 4. Discussion

The proposed method has some distinct advantages.

1. Superspecificity is for all practical purposes error-free. It relaxes the extreme demands usually made on nanopore-based sensing by reducing the burden on the detector and gives the measurement component of the method a minimalist character. As noted earlier, telling marginally different AAs apart based on the blockade requires extraordinary precision. In the methods presented this measurement complexity is considerably reduced. Thus in the 4-step procedure it is only necessary to determine in which one of 20 units a blockade has occurred. This effectively converts a high-precision analog problem into a low-precision binary one based on simple thresholding. In the 2-step procedure it is only necessary to recognize an AMP blockade in the e-cell of one of the 20 units. Again there is no need to identify an AA based on the size of the blockade it causes in the e-cell. While this requires higher measurement precision, it is much less than would be required to tell two physically similar AAs (like Leu and Ile) apart based on their blockades. No post-processing or calculations of any kind are required in either of the two procedures.
2. No information about the target protein is used or needed anywhere. This makes both procedures truly de novo in character. Currently the only practical de novo protein sequencing method is MS. However the de novo capability associated with it is the result of applying machine learning techniques to the spectral data [40], thus it is a computational feature, not an analytical one.

### Practical issues

In implementing the two methods presented here a number of practical issues may be considered. They range from severe, requiring complex solutions, to mild, with fairly straightforward solutions. In the following paragraphs some of these are discussed.

#### 1) Translocation speed control

In most nanopore-based sequencing studies the speed with which an analyte translocates through the pore usually exceeds the bandwidth of the available detector. For reviews of translocation speed control see [41,42]. Most of the methods described therein are either inapplicable or too cumbersome for the present purpose. Other methods are based on the use of organic salts [43,44] or an RTIL (room temperature ionic liquid) [45], both are aimed at increasing solution viscosity. It has also been shown that the direction of DNA translocation can be reversed at the pore entrance of an MoS_2_ pore [46]. All of these methods are limited to an increase in translocation time of about two orders of magnitude. In [47] a counterintuitive approach that uses a bilevel voltage profile is examined in theory. Thus with a positive voltage in the first segment and a negative voltage in the second segment a negatively charged analyte is induced to enter the pore and remain trapped there long enough to allow its detection with a bandwidth of about 1-10 Khz. This approach can be used even with AAs that normally have low mobility (because of low electrical charge at pH 7, such as alanine), via a change in solution pH, which increases their mobility to the required level.

#### 2) Drawing the analyte into the pore

Before an analyte can be inspected it has to be drawn into the pore from the cis chamber. In the present case 13 of the standard AAs are electrically neutral at pH 7 and the other 7 (K, R, E, D, C, H, and Y) carry a positive or negative charge. (All AAs have values that depend on the pH of the solution.) Solutions to this problem include the use of nanopipettes [48] and hydraulic pressure [49]; the latter does not depend on the charge carried by an AA.

#### 3) Redundancy in the genetic code

In the above development it has been implicitly assumed that 20 different tRNAs, one for each of the 20 standard AAs, are used to identify different AAs. As there are 61 codons coding for the 20 AAs, some AAs have multiple coding triplets and correspondingly more than one associated tRNA. As a result there are up to 61 different tRNAs that map many-to-one to the 20 RNAs. This redundancy in the genetic code can be used to increase the reliability of AA identification by using up to 61 units, each with a unique tRNA, in the parallel architecture of Figure 6. The outputs of units with different tRNAs that bind with the same AA should be the same, thus increasing the reliability of identification.

#### 6) Silo effect

In the parallel multiple unit architecture of Figure 6 every unit is like a silo: it is isolated from every other unit and works with a unique AA. As a result every unit can be tailored to the AA associated with it and designed for optimal performance based on the electrophoretic, electro-osmotic, hydrodynamic, diffusion, and other properties of that AA. For example in unit 16 in Figure S-4 in the Supplement, the only AA that can enter and pass through the pore in the e-cell to cause a blockade is Ser. Values for transmembrane voltage and hydraulic pressure can be chosen to oppose each other and optimally control the speed of translocation of Ser. Similarly the threshold used to determine if a blockade has occurred or not can be set differently in each unit based on the AA and the SNR of the nanopore in the unit’s e-cell.

### Miscellaneous notes

1. The method given here can be used for protein identification based on a small number of target residues such as described in [7] by using only the corresponding units in the parallel processor of Figure 5.
2. The cycle time is given by the time for one of the units to identify the corresponding AA. Occurrence of a blockade in the unit’s e-cell signals the end of the cycle and the start of the next.
3. In Step 4 other methods of identifying the free AA can be used in place instead of a current blockade in an e-cell. In particular this can be done with methods that are designed for bulk samples.
4. Since tRNA and AARS do not exit the cavity, a unit can be used multiple times without having to reload tRNA and AARS into it. It is also possible to design an AA identification unit with a cavity that is preloaded with a specific cognate tRNA-AARS pair.
5. Note that following deacylation in Step 3 the likelihood of reacylation of a tRNA is very low, especially since AARS needs ATP to charge tRNA and any extra ATP that might be present at the end of charging would have been filtered out in Step 2.
6. The probability of successful charging of tRNA after containment can be increased by using a tapered cavity (similar to the tapered reservoir).

## Supporting Information

One file with data tables, simulation details, notes on error analysis, and two examples of procedural traces.

