## Supplementary Information for "Label-free amino acid identification for *de novo* protein sequencing via tRNA charging and current blockade in a nanopore"

G.Sampath

---

**S-1 Geometry of reservoir-cavity structure**
**S-2 Analyte dimensions**
**S-3 Simulation of reservoir-cavity**
**S-4 Notes on the two amino acid identification procedures**
**S-5 Examples of amino acid identification with the two procedures**

Additional references in this supplement are given by numbers in square brackets starting with 50. References in the main text are numbered as listed there ([1] to [49]).

---

### S-1 Geometry of reservoir-cavity structure

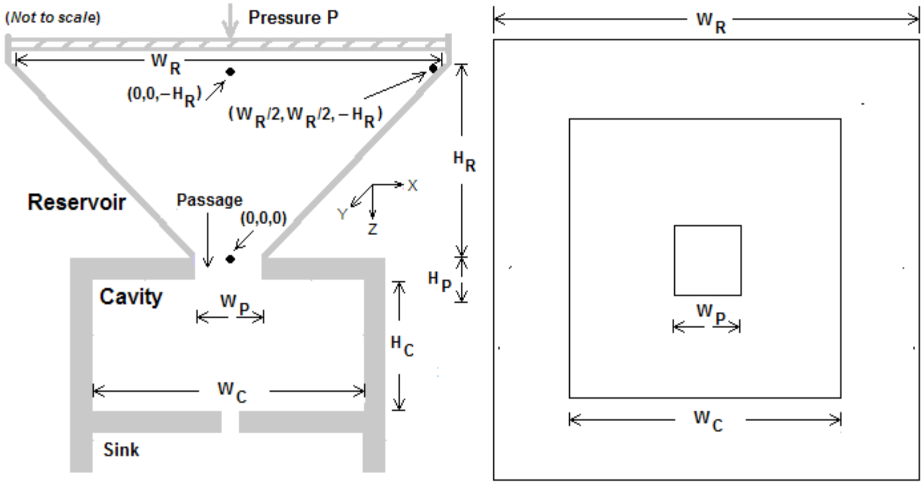


**Fig. S-1** Geometry (symmetric in X and Y) used in simulation of reservoir-cavity structure with hydraulic pressure P. Three starting points for particle are shown: (0,0,0), $(0,0,-H_R)$, and $(W_R/2, W_R/2, -H_R)$. Structure shown in plan on the right.

The reservoir is a tapered cylinder ending in a passage that leads to the cavity. The passage is a sub-micron pore. The left part of the figure appears in a slightly different form as Figure 2 in the main text.

*Dimensions of reservoir-cavity structure*

| | | |
|---|---|---|
| Reservoir: | $H_R = 10 \times 10^{-6}$ m | $W_R = 10 \times 10^{-6}$ m |
| Passage: | $H_P = 200$ nm | $W_P = 400$ nm |
| Cavity: | $H_C = 1.5 \times 10^{-6}$ m | $W_C = 8 \times 10^{-6}$ m |

These dimensions are similar to those in [27].

### S-2 Analyte dimensions

The reactants AA, tRNA, AARS, and ATP must be able to translocate from the reservoir to the cavity through the intervening passage, which is a sub-micron-sized pore (Figure S-1). Aspect ratios can be calculated for each type of analyte from a minimum volume enclosing geometric object like an ellipsoid. This was done for AMP, ATP, Phe-tRNA, Ala-AARS, Glycine, and Tryptophan with a custom R language program operating on atomic coordinate data downloaded from the RCSB website and [29]. The results are given in Table S-1 below.

**Table S-1** Dimensions of reactants involved in tRNA charging. Amino acids are represented by smallest (Glycine) and largest (Tryptophan), tRNAs by Phe-tRNA, and AARSs by Ala-AARS.

| Space or reactant | Dimensions of box/ellipsoid axes enclosing structure/reactant | | | Reactant | Axes of ellipsoid enclosing reactant | | |
|---|---|---|---|---|---|---|---|
| | Length (nm) | Breadth (nm) | Height (nm) | | Major (nm) | Minor (nm) | Minor (nm) |
| Reservoir in Figure S-1 | 10000 | 10000 | 10000 | AMP [3] | 1.67 | 0.95 | 0.4 |
| Passage in Figure S-1 | 200 | 200 | 400 | ATP [4] | 2.34 | 1.09 | 0.44 |
| Cavity in Figure S-1 | 2000 | 2000 | 1500 | Phe-tRNA [5] | 10.56 | 6.54 | 3.64 |
| Glycine (AA) [1] | 0.56 | 0.35 | 0.21 | Ala-AARS [6] | 13.94 | 9.41 | 8.46 |
| Tryptophan (AA) [2] | 1.10 | 0.7 | 0.34 | | | | |

[1,2,3,4,6] Atomic coordinate data from files.rcsb.org/ligands/view/AMP_ideal.sdf, files.rcsb.org/ligands/view/ATP_ideal.sdf, www.rcsb.org/ligand/GLY, files.rcsb.org/ligands/view/TRP_ideal.sdf, www.rcsb.org/structure/1YFS. (The last is for Ala-AARS.) [5] Atomic coordinate data from [29].

### S-3 Simulation of reservoir-cavity

The simulation procedure is similar to the procedures described in [34]. It can be viewed in the following terms:

1) Each simulation step takes place in 1 picosecond ($10^{-12}$ s).
2) A reactant (free AA, AARS, ATP, AMP, tRNA, amino-acyl tRNA) is a dimensionless particle that is reflected off the walls of the reservoir, the passage, and the cavity.
3) The particle is considered to be confined when it reaches the bottom of the cavity and stays close to it within a specified margin for a specified time.
4) The hydraulic pressure field superimposed by the piston on diffusion is assumed to result in Poiseuille flow [28] in all three segments: reservoir, passage, and cavity.
5) The displacement of a particle is equal to the sum of the diffusive translation vector and the displacement due to hydraulic pressure (which is always in the z direction).
6) In the absence of hydraulic pressure the motion of the particle is entirely due to diffusion.
7) Although hydrodynamic drag is ignored, including it in the model will not materially affect the simulation as the objective is merely to show that a particle reaches the bottom with probability 1. Its only influence would be to add a delay to the trajectory of the particle as it finds its way to the bottom of the cavity. This is because even though the hydraulic displacement is very small and is about 1/40 the size of the diffusion step it is always biased in the z direction, whereas diffusive displacement is random and isotropic and the total displacement is effectively zero in the limit. The latter is simulated as a separate case in which the pressure P is set to 0.

*Constants used*

1) Diffusion constant  D = $3.0 \times 10^{-11}$ $m^2/s$, $3.0 \times 10^{-10}$ $m^2/s$, or $3.0 \times 10^{-9}$ $m^2/s$
2) Atmospheric pressure  $P_{atm}$ = 106803 Pa (= pressure unit Pascal)
3) Viscosity of solution  0.001 Pa.sec

*Calculations*

1) The hydrodynamic velocity of a particle under Poiseuille flow is given by (Lu 2013)

$$v_{hyd} = (P_{atm} + \Delta P)\, R^2_{channel} / 8\, \eta\, H_{channel} \qquad \text{(S-1)}$$

where $P_{atm}$ = atmospheric pressure, $\Delta P$ = applied pressure, $R_{channel}$ = radius of channel (reservoir, passage, or cavity), $\eta$ = solution viscosity, and $H_{channel}$ = radius of channel.

2) The displacement of a particle is the vector sum of the displacements due to diffusion and hydraulic pressure. Diffusive movement occurs in a random direction in 3-d space and is simulated as a displacement during a small time interval $\Delta t$ (set to the simulation time step of 1 picosecond = $10^{-12}$ s). The diffusion displacement magnitude is $\sqrt{(6D\Delta t)}$, where D is the diffusion constant of the particle [34]. A random direction is generated as a unit vector on the surface of a sphere. This vector is then multiplied by the diffusion step size and the translation due to hydraulic pressure (= $v_{hyd}\, \Delta t$) added to it. The resulting position is accepted if it is inside the structure (reservoir, passage, or cavity) and rejected otherwise.
3) The *dwell time* is defined as the time continuously spent by the particle near the bottom of the cavity after reaching the bottom. The simulation is stopped if the particle dwell time exceeds $10^6$ steps.
4) The *escape ratio* is the ratio of the dwell time of the particle in the reservoir after it has reached the cavity bottom the first time to the dwell time at the bottom of the cavity.

*Simulation procedure*
The following procedure is executed with a time step of $\Delta t = 10^{-12}$ s.
**Procedure** Simulate-Particle-Trajectory
1. The particle is released at **p** = ($x_0$,$y_0$,$z_0$), a point inside the reservoir. Three representative points are used in the simulation: bottom center of reservoir, top center of reservoir, and top right corner of reservoir. Referring to the coordinate system in Figure S-1, these three points are (0,0,0), (0,0,-$H_R$), ($W_R$/2, $W_R$/2, -$H_R$). Initialize number of simulation steps to 0.
2. Generate random vector **r** to surface of unit sphere centered at current position. Multiply this by diffusion step size, which is determined by current position. The magnitude is given by $\sqrt{(6D\,\Delta t)}$, where D is the diffusion coefficient of the particle.
3. Calculate hydraulic velocity $v_{hyd}$ depending on where **p** is (R, P, or C), and hydraulic displacement in z direction as **d** = (0, 0, $v_{hyd}\,\Delta t$).
4. If **p** + **r** + **d** is in R, P, or C, set **p** = **p** + **r** + **d** and increment number of simulations by 1; else reject the move. (The latter flows from the assumption that the walls of the structure are reflective.)
5. If number of simulation steps = maximum, exit. (This maximum is $4 \times 10^8$ with pure diffusion, and $10^6$ with hydraulic pressure after the particle has reached the cavity bottom for the first time.)
7. Go to Step 2.

*Choice of parameters in simulations*
1) The value $3.0 \times 10^{-10}$ m$^2$/s for the diffusion coefficient D corresponds approximately to that for AMP [3]. The two other values used, namely $3.0 \times 10^{-11}$ m$^2$/s and $3.0 \times 10^{-9}$ m$^2$/s are representative of analytes smaller than AMP (the 20 amino acids) and larger (ATP).
2) Three values of ΔP are used: 0, 1.5, and 2.0. Anything higher than 2.0 produces considerable delays when the particle is close to the tapering wall in the reservoir. This is because the higher displacement causes the attempted move to be rejected because it falls outside and the original position retained (physically this corresponds to reflection at the wall), and this was found to happen repeatedly during the simulation. For ΔP > 0 the reservoir cavity structure used is that in Figure 4 in the main text. This allows an exit path for fluid flow out of the cavity through the pore connecting the cavity to the *cis* chamber of the e-cell.
3) The three starting points for a particle are representative of the full domain. Thus (0,0,0) and ($W_R$/2, $W_R$/2, -$H_R$) represent two extremes and (0,0,-$H_R$) an intermediate case, see Figure S-1 (almost the same as Figure 2 in the main text).

*Events monitored*
1) Time for particle to reach the bottom of the cavity: This is given by the number of steps $N_1$ taken by the particle to reach within 1 nm (= $10^{-9}$ m) of the bottom
2) Cavity entry time: This is the number of steps $N_2$ taken by the particle before it first enters the cavity.
3) Dwell time of particle near the bottom: This is given by the number of steps $N_3$ during which the particle's z coordinate is within 1 nm of the bottom

*Results*
**Table S-2** **Simulation of dimensionless particle in reservoir-cavity structure under hydraulic pressure and diffusion**

| Diffusion coefficient ($10^{-10}$ m$^2$/s) | ΔP (above atmospheric pressure) | Origin | $N_1$ = Time to enter cavity (× $10^{-6}$ s) | $N_2$ = Time to reach cavity bottom (× $10^{-6}$ s) | $N_3$ = Dwell time at bottom of cavity (× $10^{-6}$ s) | $N_4$ = Dwell time in reservoir after once reaching bottom of cavity (× $10^{-6}$ s) | Escape ratio = $N_4$ / $N_3$ |
|---|---|---|---|---|---|---|---|
| 0.3 | 0 | (0,0,0) | 1.207 | 1.376 | > 1.0 | 0 | 0 |
| | | (0,0,-$H_R$) | 1.530 | 1.698 | > 1.0 | 0 | 0 |
| | | ($W_R$/2, $W_R$/2, -$H_R$) | 1.618 | 1.786 | > 1.0 | 0 | 0 |
| 0.3 | 0.5 | (0,0,0) | 0.801 | 0.913 | > 1.0 | 0 | 0 |
| | | (0,0,-$H_R$) | 0.997 | 1.110 | > 1.0 | 0 | 0 |
| | | ($W_R$/2, $W_R$/2, -$H_R$) | 1.080 | 1.192 | > 1.0 | 0 | 0 |
| 0.3 | 1.0 | (0,0,0) | 0.587 | 0.671 | > 1.0 | 0 | 0 |
| | | (0,0,-$H_R$) | 0.753 | 0.837 | > 1.0 | 0 | 0 |
| | | ($W_R$/2, $W_R$/2, -$H_R$) | 0.826 | 0.910 | > 1.0 | 0 | 0 |
| 3.0 | 0 | (0,0,0) | 1.226 | 1.395 | > 1.0 | 0 | 0 |
| | | (0,0,-$H_R$) | 1.470 | 1.638 | > 1.0 | 0 | 0 |
| | | ($W_R$/2, $W_R$/2, -$H_R$) | 1.622 | 1.792 | > 1.0 | 0 | 0 |

| | | | | | | | |
|---|---|---|---|---|---|---|---|
| 3.0 | 0.5 | (0,0,0) | 0.783 | 0.896 | > 1.0 | 0 | 0 |
| | | $(0,0,-H_R)$ | 0.992 | 1.104 | > 1.0 | 0 | 0 |
| | | $(W_R/2, W_R/2, -H_R)$ | 1.200 | 1.312 | > 1.0 | 0 | 0 |
| 3.0 | 1.0 | (0,0,0) | 0.564 | 0.649 | > 1.0 | 0 | 0 |
| | | $(0,0,-H_R)$ | 0.743 | 0.827 | > 1.0 | 0 | 0 |
| | | $(W_R/2, W_R/2, -H_R)$ | 0.837 | 0.922 | > 1.0 | 0 | 0 |
| 30.0 | 0 | (0,0,0) | 1.176 | 1.345 | > 1.0 | 0 | 0 |
| | | $(0,0,-H_R)$ | 1.246 | 1.412 | > 1.0 | 0 | 0 |
| | | $(W_R/2, W_R/2, -H_R)$ | 1.632 | 1.801 | > 1.0 | 0 | 0 |
| 30.0 | 0.5 | (0,0,0) | 1.039 | 1.149 | > 1.0 | 0 | 0 |
| | | $(0,0,-H_R)$ | 1.272 | 1.383 | > 1.0 | 0 | 0 |
| | | $(W_R/2, W_R/2, -H_R)$ | 1.099 | 1.210\ | > 1.0 | 0 | 0 |
| 30.0 | 1.0 | (0,0,0) | 0.630 | 0.713 | > 1.0 | 0 | 0 |
| | | $(0,0,-H_R)$ | 0.633 | 0.718 | > 1.0 | 0 | 0 |
| | | $(W_R/2, W_R/2, -H_R)$ | 0.785 | 0.868 | > 1.0 | 0 | 0 |

A subset of the above table appears in the main text as the upper part of Table 1.

*Pure diffusion*
It is useful to look at what happens when there is no hydraulic pressure and the only motive force is diffusive. The following measures are used: 1) Total number of diffusive steps; 2) The final value of z; 3) The maximum value of z attained by the particle (which need not the the same as the final value of z). Table S-3 shows that with pure diffusion the particle enters the cavity in only one case, even then it does not reach the cavity bottom. In all three cases the particle is still diffusing in the reservoir at the end of the run. One can conclude that hydraulic pressure is key to the confinement of reactants to the cavity so that charging of a tRNA with a cognate AA can take place.

**Table S-3 Simulation of dimensionless particle in reservoir-cavity structure under pure diffusion**
Diffusion coefficient = $3.0 \times 10^{-10}$ $m^2/s$

| Origin | $N_5$ = Total number of steps taken ($\times 10^8$) | Value of z after $N_5$ steps ($\times 10^{-6}$ m) | Maximum z during $N_5$ steps ($\times 10^{-6}$ m) | Entered cavity? | Reached bottom of cavity? |
|---|---|---|---|---|---|
| (0,0,0) | 4 | -8.815 | 0.006 | Yes | No |
| $(0,0,-H_R)$ | 4 | -9.751 | -9.662 | No | No |
| $(W_R/2, W_R/2, -H_R)$ | 4 | -9.901 | -9.672 | No | No |

The above table appears in the main text as the lower part of Table 1.

### S-4 Notes on the two amino acid identification procedures

1) *Analyte volume*
In the main text the volume of a molecule is used as a proxy for the current blockade level (Equation 6, main text). Volume data for all three types of molecules are given in Table S-4 below.

**Table S-4** Volumes of the standard 20 amino acids, ATP, and AMP.

| **AA** | **G** | **A** | **S** | **C** | **D** | **T** | **N** | **P** | **V** | **E** | **Q** |
|---|---|---|---|---|---|---|---|---|---|---|---|
| **Mean (μ)** | 59.9 | 87.8 | 91.7 | 105.4 | 115.4 | 118.3 | 120.1 | 123.3 | 138.8 | 140.9 | 145.1 |
| **S.D. (σ)** | 2.2 | 2.3 | 1.8 | 5.0 | 2.2 | 2.3 | 4.1 | 1.8 | 3.6 | 5.3 | 5.1 |
| **AA / Other** | **H** | **M** | **I** | **L** | **K** | **R** | **F** | **Y** | **W** | **AMP** | **ATP** |
| **Mean (μ)** | 156.3 | 165.2 | 166.1 | 168 | 172.7 | 188.2 | 189.7 | 191.2 | 227.9 | 349.0 | 436.5 |
| **S.D. (σ)** | 6.1 | 1.8 | 3.4 | 4.3 | 5.9 | 9.6 | 7.4 | 8.0 | 3.8 | - | - |

Amino acid mean volumes and standard deviations are from [50]. AMP volume from Table I in [51], calculated as union of van der Waals spheres of every atom in the molecule. ATP volume approximated as 1.25 * volume of AMP

2) *Charging of tRNA with an amino acid in both procedures*

In a biological cell an amino-acyl tRNA synthetase (AARS) enzyme activates ATP and a cognate AA to form an amino acid AMP complex, charges the cognate tRNA with AA at the latter's C-terminal end and releases the AMP, then releases the charged or aminoacyl tRNA [24]. The following equations (which appear in the main text as Equations 1 through 3) summarize the process:

$$\text{AA} + \text{ATP} \rightarrow \text{Aminoacyl-AMP} + \text{pyrophosphate or PPi} \; (= P_2\,O_7^{4-}) \qquad \text{(S-2)}$$
$$\text{Aminoacyl-AMP} + \text{tRNA} \rightarrow \text{Aminoacyl-tRNA} + \text{AMP} \qquad \text{(S-3)}$$
$$P_2\,O_7^{4-} + H_2O \rightarrow 2\,HPO_4 \;\; + 2\,H^+ \qquad \text{(S-4)}$$

This process can be used *in vitro* to charge a tRNA with a free AA. When the latter is exposed to distinct tRNAs, their cognate AARSs, and ATP in solution, one of the tRNAs will get charged with AA. This may be done in parallel with N ($20 \le N \le 61$) AA molecules with N different tRNAs. Here 20 is the number of standard AAs, and 61 the number of coding triplets in the genetic code. The latter is due to 'wobble' in the third position of the triplet [24].

3) *Identifying an AA from an AMP blockade in the two-step procedure*

Of the three analyte types, the 20 amino acids have the lowest volumes. The volume of AMP ($349.0 \times 10^{-3}$ nm$^3$) is at least 50% higher than that of the largest amino acid (W, Tryptophan, volume = $227.9 \times 10^{-3}$ nm$^3$). The volume ratio between AMP and ATP is 1:1.25. Therefore the blockade due to AMP can be easily distinguished from the blockade due to any AA, and sufficiently reliably from that due to ATP, by setting two thresholds:

$$Th_{low} = (\mu_W + \mu_{AMP})/2 \qquad \text{(S-5a)}$$

and

$$Th_{high} = (\mu_{AMP} + \mu_{ATP})/2 \qquad \text{(S-5b)}$$

Additionally a lower threshold of $Th_{noise}$ is set to distinguish the blockade due to G from baseline noise (see Item 5 below). This leads to

$$Th_{low} < V_{AN} < Th_{high} \quad \rightarrow \quad AN = AMP \qquad \text{(S-6a)}$$
$$Th_{noise} < V_{AN} < Th_{low} \quad \rightarrow \quad AN = AA \qquad \text{(S-6b)}$$
$$V_{AN} > Th_{high} \quad \rightarrow \quad AN = ATP \qquad \text{(S-6c)}$$

*4) Error analysis in the two-step procedure*

In analyzing the two-step procedure for potential errors the following assumptions are made:

1) Superspecificity applies *in vitro* so that charging of a tRNA in the presence of a cognate AA always occurs and there are no charging errors. Correction of errors by AARS (Bergtom 2018), as occurs *in vivo*, is not considered here.

2) If a tRNA and its cognate AA are not present then the AARS cognate to the tRNA does not activate ATP, which means that there is no AMP release (Equations S-2 and S-3).

The key step in the identification procedure is determining if the tRNA is charged or not. This determination depends only on the current blockade levels observed in the e-cell. With perfect identification the following two conditions hold:

a) The blockade level of AMP can always be distinguished from those due to any and all of the ATP molecules as well as due to any AA; and

b) When the tRNA is charged the released AMP causes a blockade of characteristic size.

In practice identification may not be perfect. Errors can be divided into *false positives* and *false negatives*. A false positive occurs when the tRNA is not charged but the output reads 'charged', while a false negative occurs when the tRNA is charged but the output reads 'not charged'. The probabilities of these two occurrences can be estimated by using $V_{AN}$ as a proxy for the measured current blockade $I_{AN}$ (Equation 6 in the main text). Such analyses are usually simplified by assuming the relevant variables to be normally distributed with some mean μ and standard deviation σ.

Let $F_X(x, \mu, \sigma)$ be the cumulative distribution function (cdf) of the gaussian variable X, the current blockade level for analyte X. With AN = AA, ATP, or AMP, an observed blockade is evidence of translocation of an amino acid AA if

$$\mu_{AA} - k_{AA}\,\sigma_{AA} \; < \; V_{AA} \; < \; \mu_{AA} + k_{AA}\,\sigma_{AA} \qquad \text{(S-7)}$$

and of ATP (AMP) if

$$\mu_{ATP(AMP)} - k_{ATP(AMP)}\,\sigma_{ATP(AMP)} \; < \; V_{ATP(AMP)} \qquad \text{(S-8)}$$

for some positive integer value k times the spread of the distribution of analyte volume. The probabilities of false negatives and false positives are calculated next. The calculations are on the conservative side and use a spread of 5σ (k = 5) about the mean of the normal distributions involved.

(i) *False negatives*
When a tRNA is charged, the tRNA is bound to the AA, and an ATP molecule breaks down into AMP and Ppi molecules. The AMP molecule translocates to *trans*. However the output signal reads 'not charged'. This misread could result from the AMP causing an AA-sized blockade. The probability of this is

$$P_{FN\text{-}AA} = F_{AMP}(Th_{low}, \mu_{AMP}, \sigma_{AMP}) - F_{AMP}(\mu_{AA} - 5\,\sigma_{AA}, \mu_{AMP}, \sigma_{AMP}) \qquad \text{(S-9)}$$

This leads to Figure 8A in the main text.
(ii) *False positives*
In this case the tRNA is not charged, which means that there is a free AA and no reduction in the number of ATP molecules. However the signal says 'charged'. In this misread signal it would appear as if an AMP translocated to *trans* (along with other free ATP molecules in the solution), or possibly one of the ATP looks like an AMP and the AMP looks like an ATP. The latter has a much smaller probability and need not be considered. The probability of this is

$$P_{FP\text{-}AA} = F_{AA}(Th_{high}, \mu_{AA}, \sigma_{AA}) - F_{AA}(Th_{low}, \mu_{AA}, \sigma_{AA}) \qquad \text{(S-10)}$$

This leads to Figure 8B in the main text.
(iii) *False negatives and positives averaged over all amino acids in a proteome*
The standard 20 amino acids occur with different frequencies in the proteome of an organism. Figure S-2 shows their frequency distribution in the human proteome (Uniprot Id UP000005640; 11326153 residues).

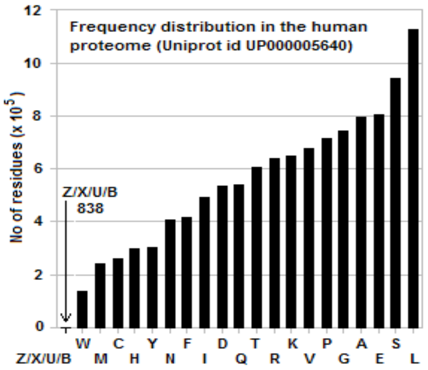


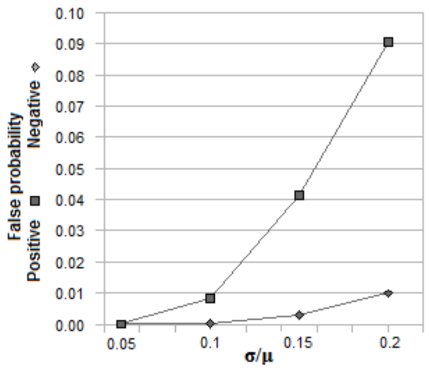


**Figure S-2 (left)** Frequency distribution of standard amino acids in human proteome (Uniprot Id UP000005640)
Proteome has 11326153 residues, including 838 residues with non-standard one-letter abbreviations (Z, X, U, and B).

**Figure S-3 (right)** Probability of false negatives and of false positives over the full set of proteins in the human proteome (Uniprot Id UP000005640) weighted by frequency of occurrence of each amino acid. (See Figure S-2 for frequency distribution of the 20 amino acids in the human proteome.)

If the two types of errors are averaged over all 20 amino acids weighted by their frequency of occurrence in the human proteome, the result is Figure S-3, which appears in the main text as Figure 8C. In this case the maximum weighted error probability is 9.5% for σ = 0.2μ.

### S-5 Examples of identification of AA = Ser with the two procedures

Figures S-4 and S-5 trace through the two AA identification procedures using AA = Ser.

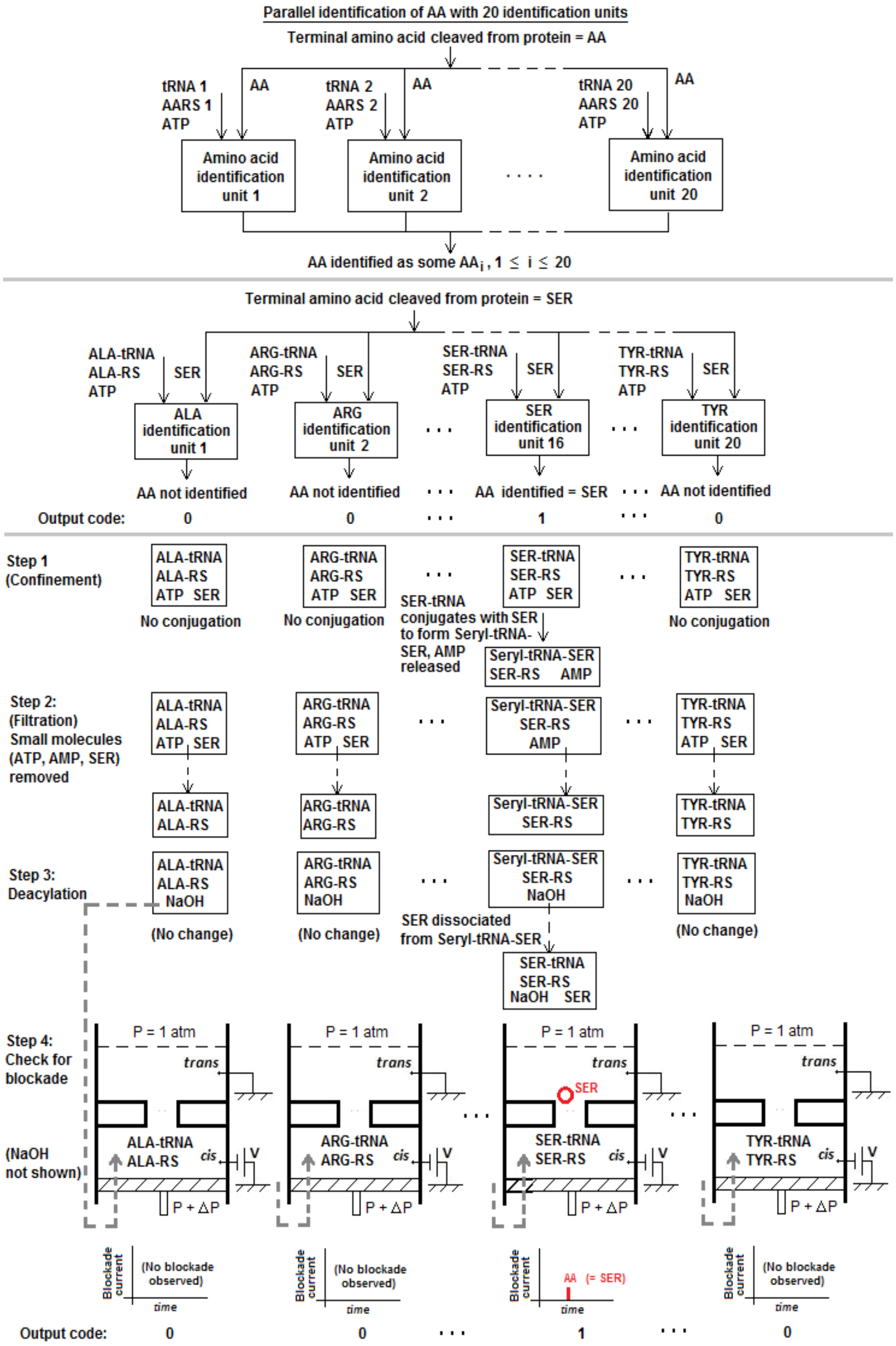


**Fig. S-5** Trace through 4-step procedure for AA = Ser

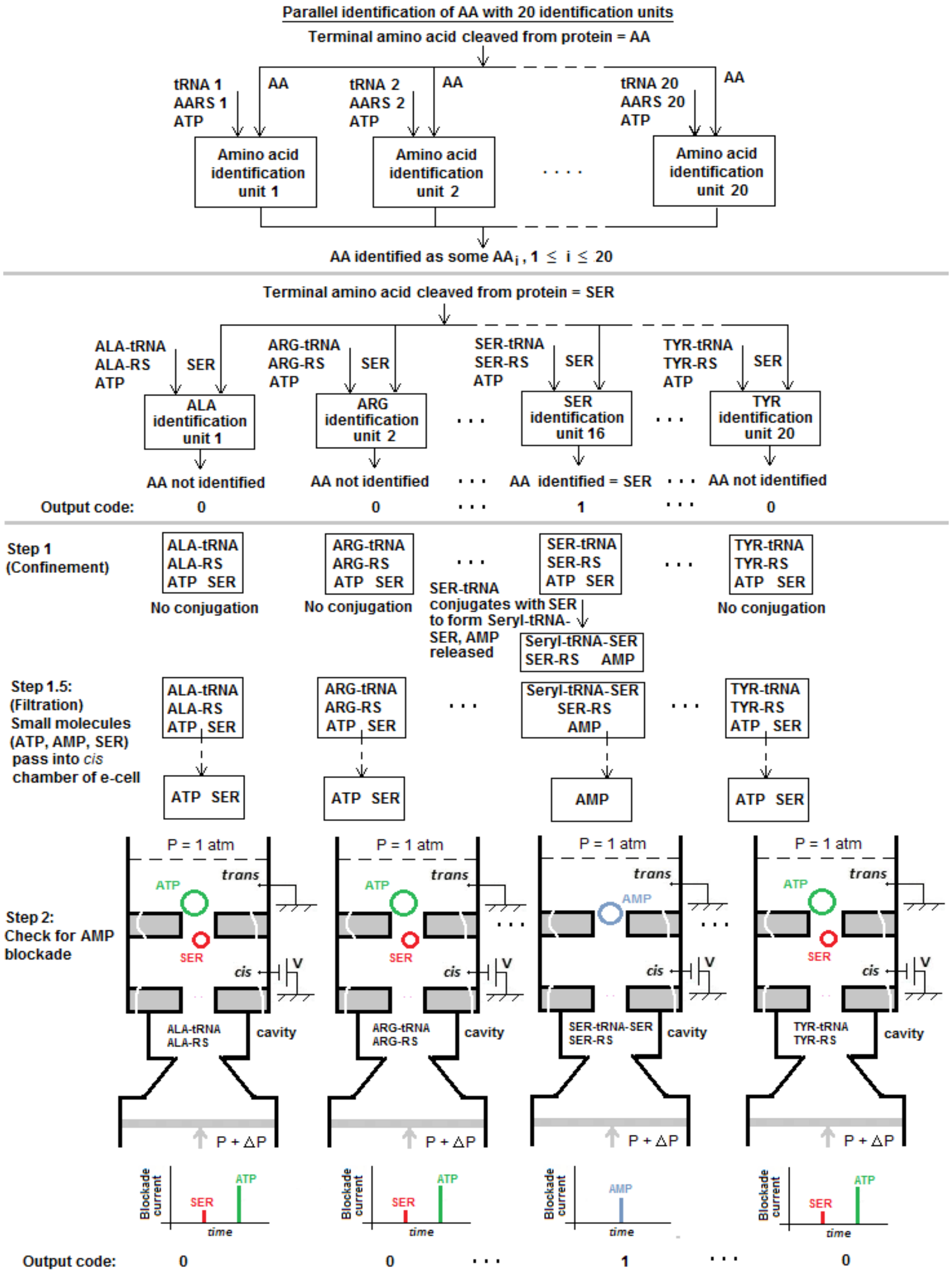


**Fig. S-6** Trace through 2-step procedure for AA = Ser
